# MetaRiPPquest: A Peptidogenomics Approach for the Discovery of Ribosomally Synthesized and Post-translationally Modified Peptides

**DOI:** 10.1101/227504

**Authors:** Hosein Mohimani, Alexey Gurevich, Kelsey L. Alexander, C. Benjamin Naman, Tiago Leão, Evgenia Glukhov, Nathan A. Moss, Tal Luzzatto-Knaan, Fernando Vargas, Louis-Felix Nothias, Nitin K. Singh, Jon G. Sanders, Rodolfo A. S. Benitez, Luke R. Thompson, Md-Nafiz Hamid, James T. Morton, Alla Mikheenko, Alexander Shlemov, Anton Korobeynikov, Iddo Friedberg, Rob Knight, Kasthuri Venkateswaran, William Gerwick, Lena Gerwick, Pieter C. Dorrestein, Pavel A. Pevzner

## Abstract

Ribosomally synthesized and post-translationally modified peptides (RiPPs) are an important class of natural products that include many antibiotics and a variety of other bioactive compounds. While recent breakthroughs in RiPP discovery raised the challenge of developing new algorithms for their analysis, peptidogenomic-based identification of RiPPs by combining genome/metagenome mining with analysis of tandem mass spectra remains an open problem. We present here MetaRiPPquest, a software tool for addressing this challenge that is compatible with large-scale screening platforms for natural product discovery. After searching millions of spectra in the Global Natural Products Social (GNPS) molecular networking infrastructure against just six genomic and metagenomic datasets, MetaRiPPquest identified 27 known and discovered 5 novel RiPP natural products.

## Introduction

Natural products are at the center of attention as new pharmaceutical leads, as exemplified by the recent discoveries of novel classes of bioactive natural product drugs^1-4^. Complementing this, the recent launch of the Global Natural Products Social (GNPS) molecular networking infrastructure^5^ brought together over a thousand laboratories worldwide that have already generated an unprecedented amount of tandem mass spectra of natural products. However, to transform natural product discovery into a high-throughput technology and to fully realize the promise of the GNPS project, new algorithms are needed for natural products discovery^6-10^. Indeed, while spectra in the GNPS molecular network represent a gold mine for future chemical discoveries, their interpretation remains a bottleneck due to the large volume of data produced by modern mass spectrometers and unavailability of computational platforms for data processing.

The efforts present herein focus on *Ribosomally synthesized and Post-translationally modified Peptides (RiPPs)*, a rapidly expanding group of natural products with applications in pharmaceutical and food industries^11^. RiPPs are produced by *RiPP Synthetases (RiPPS)* through the *Post Ribosomal Peptide Synthesis (PRPS)* pathway^11^. RiPPs are initially synthesized as *precursor peptides*, encoded by RiPP *structural genes*. The RiPP structural genes are often quite short, making their annotation difficult^12^. A precursor peptide consists of a prefix *leader peptide* appended to a suffix *core peptide*. A leader peptide is important for recognition by the *RiPP post-translational modification enzymes* and for exporting the RiPP out of the cell. The core peptide is post-translationally modified by the RiPP biosynthetic machinery, proteolytically cleaved from the leader peptide to yield the *mature RiPP*, and exported out of the cell by transporters. The precursor peptide and the enzymes responsible for post-translational modifications (PTMs), proteolytic cleavage, and transportation usually appear in a contiguous *biosynthetic gene cluster (BGC)* of a RiPP within a microbial genome. The length of the microbial RiPP-encoding BGCs typically varies from 1,000 to 40,000 bp (average length 10,000 bp), larger than the current length of short reads generated by next generation sequencing (350bp), and making DNA assembly a critical part of any short read based RiPP discovery method.

Genome mining refers to the informatics-based structural interpretation of a natural product BGC to infer information about the natural product itself. The discoveries of coelichelin in *Streptomyces coelicolor*^13,14^ and orfamide in *Pseudomonas fluorescens* Pf-5^14b,14c^ were the first examples of genome mining^13,14^ that were followed by discoveries of various bioactive RiPPs in microbial samples. Donia et al.^15^ discovered lactocillin, a thiopeptide antibiotic from human vaginal isolates, that showed activity against vaginal pathogens. Zhao et al.^16^ discovered eight novel lanthipeptides with antibiotic activity from a ruminant bacterium. Freeman et al. and Wilson et al.^17,18^ used metagenome mining of a sponge to assign a BGC to the known RiPP polytheonamide, with post-translational modifications distributed across 49 residues. Thus, large-scale metagenomics projects, such as Earth Microbiome Project^19,19b^, American Gut Project^20^, and Human Microbiome Project^21,22,22b^, have the potential to contribute to RiPP discovery, provided that improved bioinformatics tools for the enhanced identification of novel RiPPs are available. However, discovery of lactocillin and other recently identified RiPPs were not achieved by an automated process, but rather used time-consuming manual technologies that required information about individual (isolated) bacterial genomes. Our goal is to discover RiPPs directly from metagenomic information using a fully automated approach.

While recent analysis of thousands of bacterial and fungal genomes has already resulted in the discovery of many putative BGCs, including *ca*. 20,000 RiPP-encoding BGCs in the Integrated Microbial Genome Atlas of Biosynthetic Gene Clusters (IMG-ABC), connecting these BGCs to their metabolites has not kept pace with the speed of microbial genome sequencing^23^. Currently, only 35 out of these roughly 20,000 RiPP-encoding BGCs in IMG-ABC have been experimentally connected to their RiPPs^23,24^. Linking this impressive number of RiPP-encoding BGCs to unknown RiPPs requires the development of novel computational tools.

Kersten et al.^25^ introduced the peptidogenomics approach to RiPP discovery, which refers to finding sequential amino acid tags from tandem mass spectra (peptidomics) and mining them in assembled DNA reads obtained from the same sample. Mohimani et al.^12^ introduced RiPPquest, the first automated approach to RiPP discovery by mass spectrometry and genome mining. This tool is based on *Peptide-Spectrum Matches (PSMs)*, which are generated by aligning predicted spectra of putative RiPPs discovered by genome mining to mass spectra. If a *PSM* between a candidate RiPP from the assembled genome and a spectrum is statistically significant, then RiPPquest reports it as a putative annotation of the spectrum. RiPPquest resulted in identification of the lanthipeptide ‘informatipeptin’, the first natural product discovered in a fully automatic fashion by a computer. However, RiPPquest has a number of limitations: (a) RiPPquest is limited to lanthipeptides which constitutes only one of 19 classes of RiPPs^11^, (b) RiPPquest is designed for small genomes and small spectral datasets, making it rather slow in the case of large metagenomic datasets and the entire GNPS infrastructure, (c) RiPPquest does not report the statistical significance of identified RiPPs, the key requirement for any high-throughput peptide identification tool, and (d) RiPPquest is limited to searches for a predefined set of post-translational modification (PTMs) and does not enable blind searches for new PTMs.

Here we present MetaRiPPquest for high-throughput RiPP discovery that improves on RiPPquest in several critical aspects: (a) MetaRiPPquest identifies several important classes of lanthipeptides, lassopeptides, linear azole containing peptides (LAPs), linaridins, glycocins, cyanobactins, and proteusins (versus only lanthipeptides by RiPPquest), (b) algorithmic improvements in MetaRiPPquest increased speed by two orders of magnitude compared to RiPPquest, thus enabling searches of the entire GNPS database against metagenomes, (c) MetaRiPPquest implements a new approach for estimating statistical significance of identified PSMs, (d) MetaRiPPquest is capable of searching for RiPPs with unusual modifications in a blind mode, (e) MetaRiPPquest, in contrast to RiPPquest (which was designed for analyzing low-resolution spectra), utilizes the power of high-resolution mass spectrometry, and (f) MetaRiPPquest features a web-based interface through the GNPS infrastructure.

## Results

### Brief description of MetaRiPPquest

Figure 1 shows the MetaRiPPquest pipeline, which includes the following steps (see Methods section): (a) selecting a biological sample (an isolated microbe or a bacterial/fungal community), (b) generating DNA sequence data, (c) assembly of DNA sequence data with SPAdes^26^ or MetaSPAdes^27^, (d) identifying putative RiPP precursor peptides using antiSMASH^28^ and BOA^28a^ (<u>B</u>acteriocin gene block and <u>O</u>peron <u>A</u>ssociator), and constructing a database of putative RiPP structures (as well as a decoy database), (e) generating tandem mass spectra for samples, (f) matching spectra against the constructed RiPP structure database using Dereplicator^29^, and (g) enlarging the set of described RiPPs via spectral networking^30,31^.

**Figure 1.**
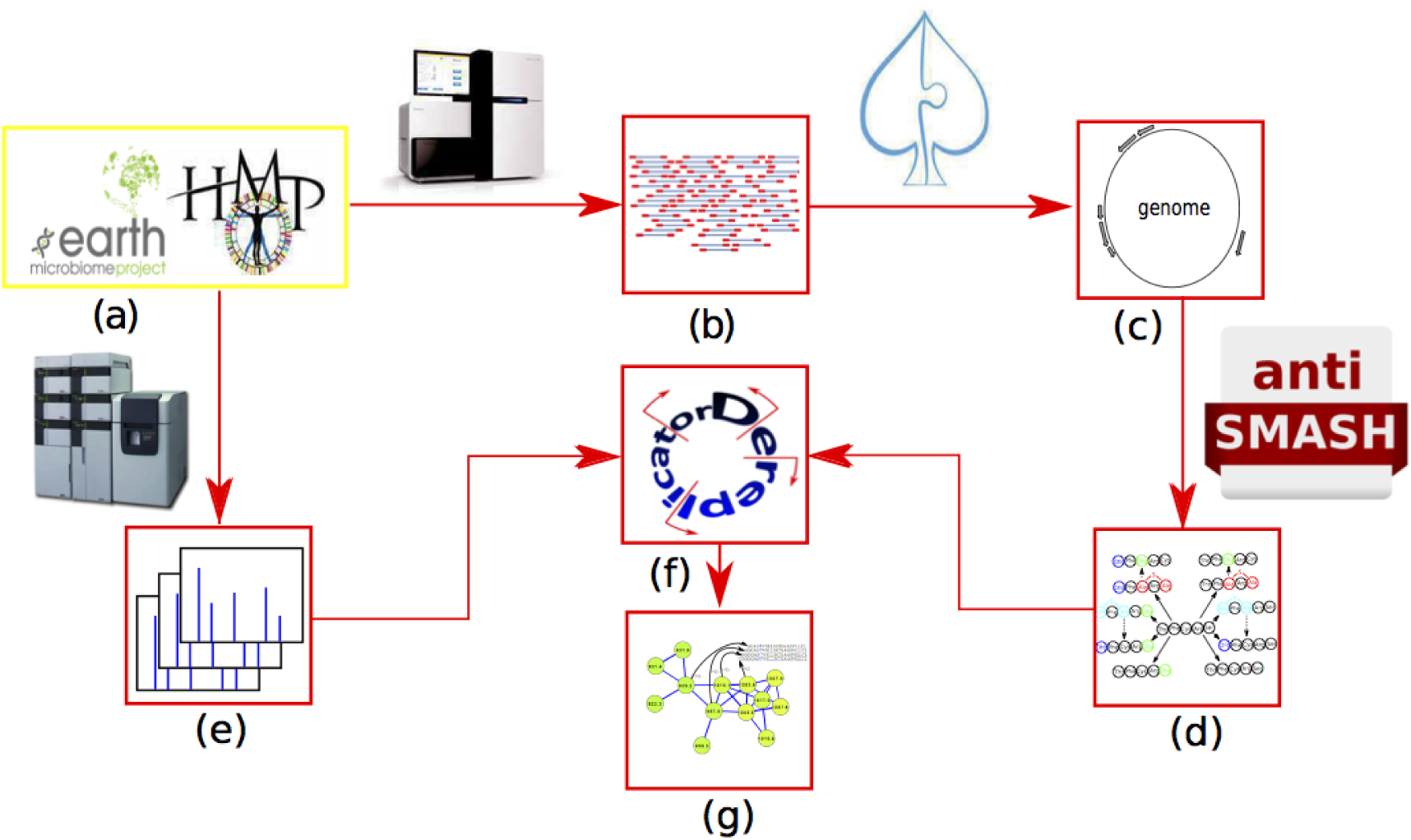
The MetaRiPPQuest pipeline includes the following steps: (a) selecting a biological sample (an isolated microbe or a bacterial/fungal community), (b) generating DNA sequence data, (c) assembly of DNA sequence data with SPAdes^26^ or metaSPAdes^27^, (d) identifying putative RiPP precursor peptides using antiSMASH^28^ and constructing a database of putative RiPPs structures (as well as a decoy database), (e) generating tandem mass spectra, (f) matching spectra against the constructed RiPP structure database using Dereplicator^29^, and (g) enlarging the set of described RiPPs via spectral networking^30,31^.

### Datasets

We analyzed the following paired datasets of spectra and genome/metagenome data (all datasets, with the exception of the BACIL dataset, contain high-resolution spectra):

- *Standard dataset (STANDARD).* This small dataset consists of 18 spectra of known RiPPs that were used for benchmarking MetaRiPPquest (GNPS datasets MSV000079506 and MSV000079622). Spectra were collected from purified RiPPs from *Prochlorococcus marinus* MIT 9313 (four analogs of prochlorosins), *Geobacillus thermodenitrificans* NG80 (geobacillin), *Bacillus subtilis* NCIB 3610 (sublancin), *Bacillus halodurans* C-125 (haloduracin), *Lactococcus lactis* (lacticin), *Bacillus cereus* SJ1 (two analogs of bicereusins) and *Ruminococcus flavefaciens* FD-1 (eight analogs of flavecins). For these strains, we used genome sequence information available from the NCBI RefSeq database. AntiSMASH identified 70 BGCs in these genomes, including 29 RiPP-encoding BGCs. Since the genome sequence of the lacticin producer is not available, we searched its spectrum against its described biosynthetic gene cluster ^31x^.
- *Actinomycetes dataset (ACTI).* This dataset consists of 473,135 spectra from bacterial extracts of 36 *Actinomycetales* strains with sequenced genomes^32,33^ (GNPS datasets MSV000078839 and MSV000078604). We downloaded sequence information for these 36 genomes from the NCBI RefSeq database. AntiSMASH identified 1,140 BGCs in these genomes, including 168 RiPP-encoding BGCs. Furthermore, we downloaded and mixed the short reads from 21 out of 36 strains that were available from the NCBI Short Reads Archive (read length 150 bp, insert sizes varying between 200bp to 300bp). We randomly down-sampled each dataset to 10 million reads (resulting in an approximate 300-fold coverage), and mixed all the reads to simulate a metagenomic dataset for this sample. Running MetaRiPPquest on the separate genomes from this dataset resulted in the same set of identified RiPPs as obtained from the simulated metagenome.
- *Bacillus dataset (BACIL).* This dataset consists of 40,051 low-resolution spectra from bacterial extracts of two *Bacillus* strains with known genomes^34^ (MSV000078552). We downloaded genome sequence information for these isolates from the NCBI RefSeq dataset.
- *Space station dataset (SPACE).* This dataset consists of 58,422 spectra from bacterial extracts of 21 isolated strains from the international space station (MSV000080102). Among these strains, twelve are *Staphylococcus*, six *Bacillus*, four *Enterobacteria* and one *Acinetobacter* strain. The complete genomes are available for all of these strains.^35,36^
- *Sponge dataset (SPONGE).* This dataset contains 223,135 spectra from bacterial extracts of *Theonella swinhoei* (GNPS dataset MSV000078670). Wilson et al.^18^ used the SPONGE dataset to analyze for the RiPP polytheonamide. We searched spectra from the SPONGE dataset against the genome of *Theonella swinhoei* symbiont *Candidatus Entotheonella* sp. TSY1. AntiSMASH identified 27 BGCs in the symbiont genome, including one RiPP-encoding BGC.
- *Cyanobacteria dataset (CYANO).* This dataset consists of 11,921,457 spectra from the extracts of 317 cyanobacterial samples^37^ (GNPS dataset MSV000078568). Each sample represents a mini-metagenome^27,38^ with one or a few highly abundant strains. The metagenomic reads were collected from 195 of these samples.

### Genome mining

MetaRiPPquest uses antiSMASH and BOA for identification of RiPP-encoding BGCs and has two genome mining modes for selecting Open Reading Frames (ORFs), a slow all-ORF mode introduced in RiPPquest^12^, and a new fast motif-ORF mode. The all-ORF approach analyzes all short ORFs within a BGC, while the motif-ORF approach relies on RiPP motif finding^39^ to narrow the set of putative RiPP-encoding ORFs.

We illustrate positive and negative features of these approaches through genome mining of the *Streptomyces roseosporous* NRRL 11379 genome obtained from the ACTI dataset. AntiSMASH found 30 BGCs in this genome, including six RiPP-encoding BGCs in this genome. Within these six BGCs, the motif-ORF approach identified only two short ORFs matching core RiPP motifs, while the all-ORF approach identified 14,694 short ORFs. When analyzing all 36 of the ACTI strains, antiSMASH discovered 1,140 BGCs, including 168 RiPP-encoding BGCs. MetaRiPPquest in the motif-ORF and all-ORF modes identified 67 and 565,138 short ORFs, respectively. This example illustrates that the motif-ORF mode may result in a four order of magnitude reduction in the number of ORF candidates as compared to the all-ORF mode. However, antiSMASH predictions are based on searching for a set of known motifs, therefore the motif-ORF mode misses some ORFs with novel RiPP motifs. BOA is based on identifying known proximal genes (“context genes”) that reside next to the RiPP, rather than by the RiPP sequence itself. In that manner, BOA identifies non-orthologous RiPP replacements if those RiPPs maintain homologous context genes. However, if the RiPPs do not have context genes, BOA may not detect those RiPPs. Also, since BOA is trained on bacteriocin context genes only, it is especially suited for that type of RiPPs.

Although the all-ORF mode searches a larger set of ORFs than the motif-ORF mode, it does not necessarily result in an increased number of identified RiPPs after matching ORFs against the spectral dataset. Indeed, the PSMs that are statistically significant in the motif-ORF mode may become statistically insignificant in the all-ORF mode because the search space in the all-ORF mode is orders of magnitude larger than in the motif-ORF mode resulting in an increased false discovery rate (FDR). Because MetaRiPPquest only reports statistically significant PSMs, the all-ORF mode may miss some peptides identified in the motif-ORF mode. Conversely, because MetaRiPPquest searches more ORFs in the all-ORF mode than in the motif-ORF mode, the motif-ORF mode may miss some peptides identified in the all-ORF mode.

### RiPP identification

Below we describe applications of MetaRiPPquest to various datasets:

#### STANDARD

For STANDARD dataset in the all-ORF mode, MetaRiPPquest identified 18 RiPPs including prochlorosin^40^, geobacillin^41^, sublancin^42^, haloduracin^43^, lacticin 481^44^, bicereucin^45^, flavecin^16^ with p-values ranging from 8•10^−14^ to 5•10^−55^ (Table 1). In contrast, MetaRiPPquest in the motif-ORF mode identified only five out of 18 RiPPs since antiSMASH failed to predict 13 out of 18 RiPP-encoding ORFs.

**Table 1:**
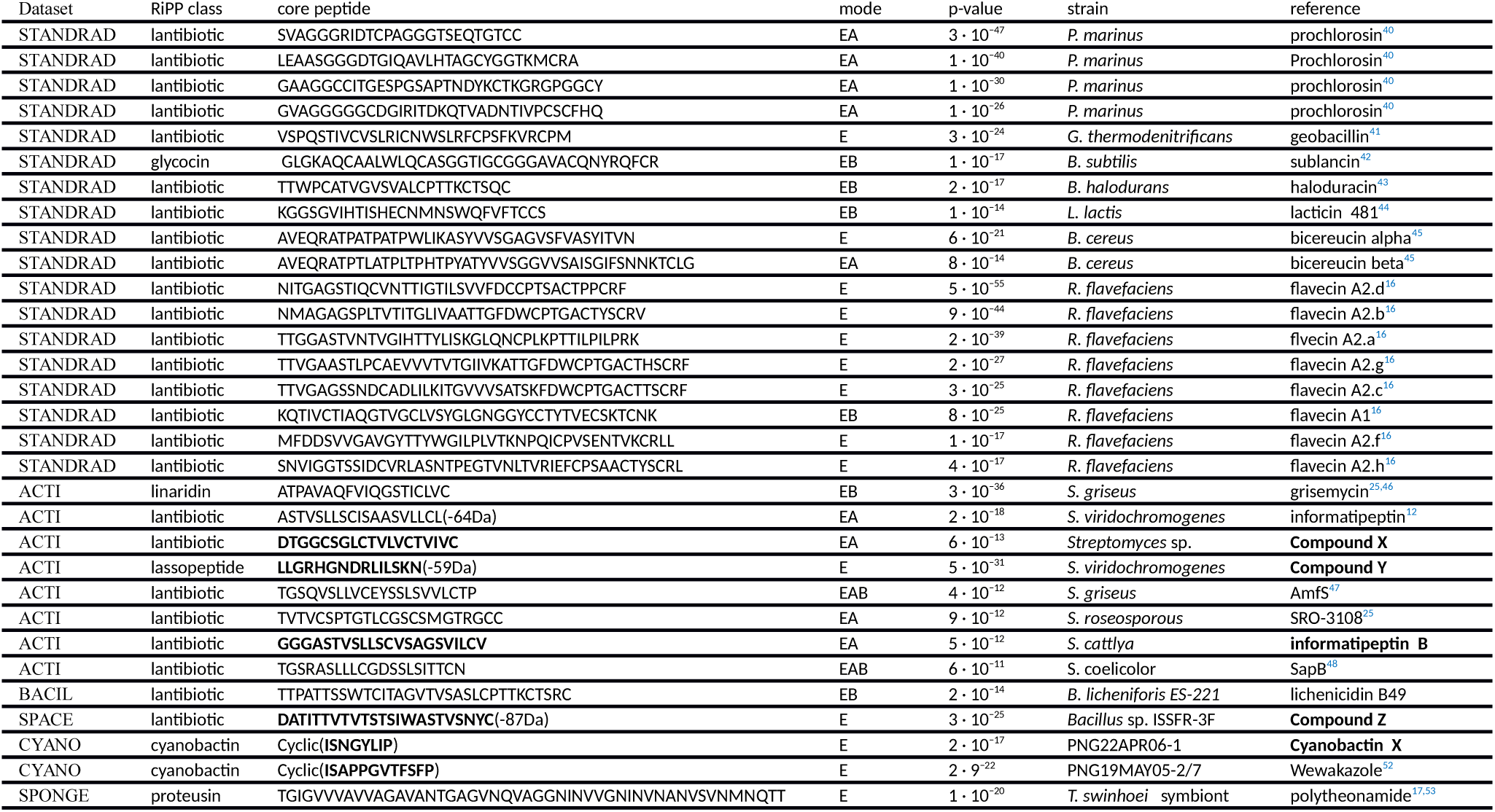
MetaRiPPquest identified 26 known and five novel RiPPs in STANDARD, BACIL, SPACE, ACTI, and SPONGE datasets. All identified RiPPs are linear except for cyclic cyanobactins. The novel RiPPs are shown in bold. The mode stands for the mode of discovery. A stands for RiPPs discovered by antiSMASH motif search, B stands for RiPPs discovered by BOA search, and E stands for RiPPs discovered by exhaustive search. Overall, 11 out of 31 RiPPs are predicted by antiSMASH motif search, 8 out of them by BOA search, and 17 only with exhaustive search.

#### ACTI

Figure 2 shows a comparison of performance of MetaRiPPquest with all-ORF and motif-ORF genome mining approaches on the ACTI dataset. At the extremely conservative 0% FDR, MetaRiPPquest in the motif-ORF mode identified three novel RiPPs and five known RiPPs. The five known RiPPs include the linaridin grisemycin at p-value of 3•10^−36^ (from *Streptomyces griseus* IFO 13350^25,46^), lantibiotics AmfS and SRO-3108 at p-value of 4•10^−12^ and 9•10^−12^ (from *Streptomyces* roseosporous^25,47^), lantibiotic informatipeptin at p-value 5•10^−12^ (from *Streptomyces viridochromogenes*^12^), and the lantibiotic SapB at p-value of 4•10^−12^ (from *Streptomyces coelicolor* A3(2)^48^). The three novel RiPPs include a class II lantibiotic (referred to as Compound X) from *Streptomyces sp.* CNT360 with p-value 6•10^−13^, an informatipeptin-like lantibiotic (referred to as informatipeptin B) from *Streptomyces cattleya* with p-value 5•10^−12^, and a lassopeptide (refered to as Compound Y) from *Streptomyces viridochromogenes* with p-value 5•10^−31^. MetaRiPPquest in the all-ORF mode identified only two known RiPPs at 0% FDR (grisemycin and AmfS). Note that while the all-ORF mode improves on the motif-ORF mode for the STANDARD dataset, the motif-ORF mode improves on the all-ORF mode for the ACTI dataset. MetaSPAdes assembled the simulated ACTI metagenome into 6,204 contigs with lengths greater than 1000 bp, with total length of 99.9 Mb and N50 of around 60 kb. The longest contig was approximately 756 kb. AntiSMASH identified 353 BGCs (66 RiPPs) in the assembled metagenome. All of the identified RiPPs were re-identified using the simulated metagenome rather than individual genomes.

**Figure 2.**
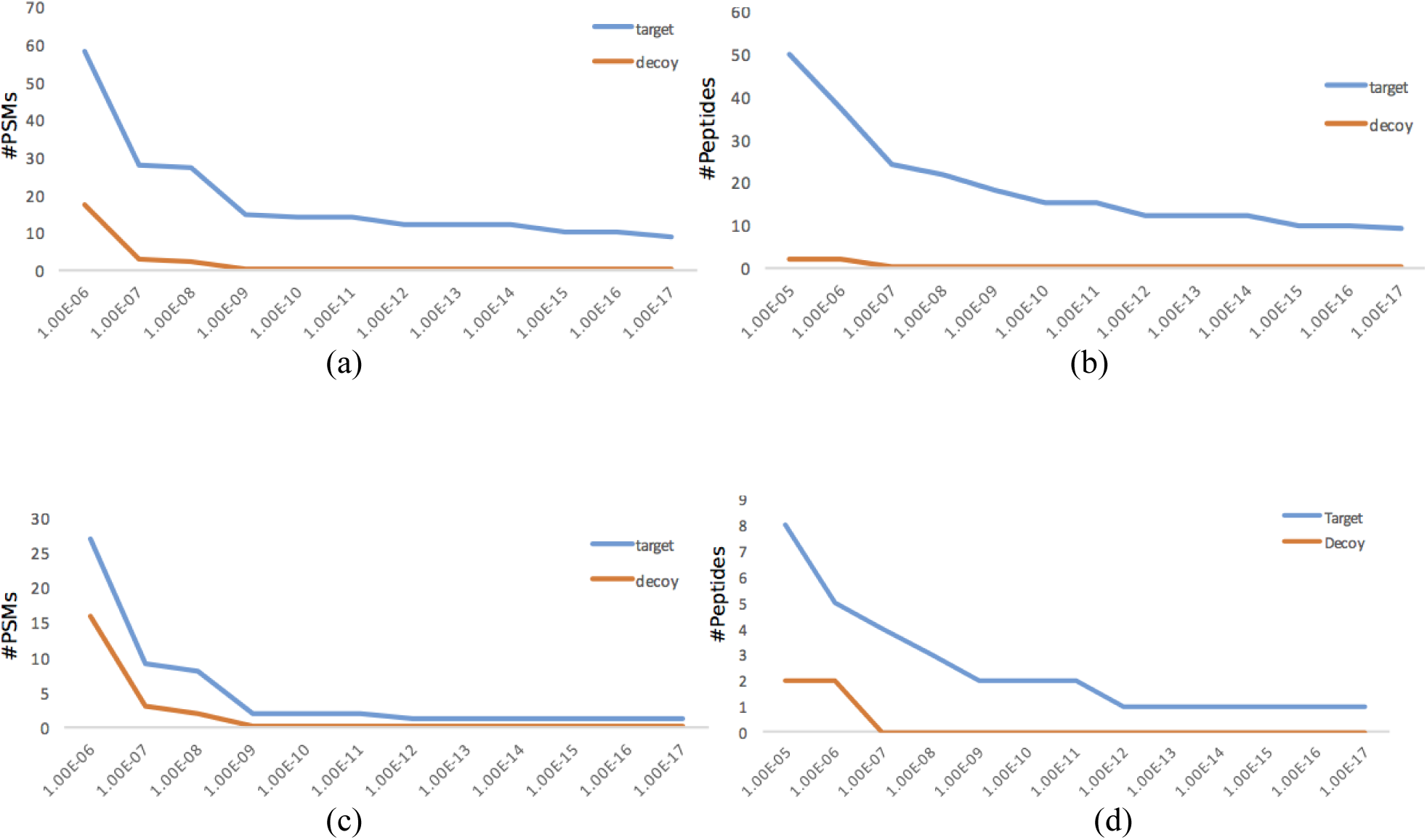
The histograms of p-values of PSMs/Peptides identified by MetaRiPPquest for the target and decoy databases in the search of ACTI dataset in the all-ORF (a,b) and motif-ORF (c,d) modes.

#### BACIL

AntiSMASH identified 12 BGCs (one RiPP-encoding BGCs) in *B. amyloliquefaciens* FZB42 and 11 BGCs (four RiPP-encoding BGCs) in *B. licheniformis* ES-221. MetaRiPPquest identified two known RiPPs from the BACIL dataset. Lichenicidin A, a class II lantibiotic^51^, was identified from a novel producer *Bacillus licheniformis* ES-221. Plantzocilin A, a linear azole containing peptide^49,50^, was identified from the known producer *Bacillus amyloliquefaciens* FZB42.

#### SPACE

AntiSMASH identified 119 BGCs, including 27 RiPP-encoding BGCs, in this dataset. MetaRiPPquest in the all-ORF mode identified one novel lantibiotic named ‘compound Z’ from three *Bacillus* strains (*Bacillus* sp. ISSFR-3F, *Bacillus* sp. S1-R2-T1 and *Bacillus* sp. S1-R3-J1).

#### SPONGE

AntiSMASH identified 27 BGCs (including four RiPP-encoding BGCs) in the SPONGE metagenome. MetaRiPPquest identified the known proteusin RiPP polytheonamide^17,52^ with a p-value 10^−20^ encoded by one of these four BGCs.

#### CYANO

AntiSMASH identified 2,898 BGCs in the 195 cyanobacterial metagenomes, including 491 RiPP-encoding BGCs. MetaRiPPquest identified the known RiPP wewakazole^53^ and a novel RiPP named cyanobactin X in all-ORF mode.

### Novel RiPPs

Below we describe five novel RiPPs and wewakazole, a known RiPP with a novel gene cluster, identified by MetaRiPPquest.

Informatipeptin B (NGGGASTVSLLSCVSAGSVILCV) is a novel lantibiotic identified with a p-value of 5•10^−12^ (Figure 3) in the ACTI dataset. Informatipeptin B differs from informatipeptin^12^ in only 5 amino acids (shown in bold). The most abundant analog of informatipeptin B (mass of 1870.95 Da) corresponds to six dehydrations (unmodified peptide mass of 1979.01). The BGC of informatipeptin B is similar to the BGCs encoding class III lantibiotics (AmfS and SapB). Three out of 10 nodes in the spectral network of informatipeptin B have been identified as its analogs: informatipeptin B1 (addition of N-terminal amino acid N), informatipeptin B2 (truncation by one amino acid), and informatipeptin B3 (truncation by two amino acids). These analogs of informatipeptin B (with stepwise N-terminal leader processing) provide additional evidence that informatipeptin B is a novel RiPP. Four other nodes in the spectral network have been identified as sodium and potassium adducts (+22 Da and +38 Da mass shifts).

**Figure 3.**
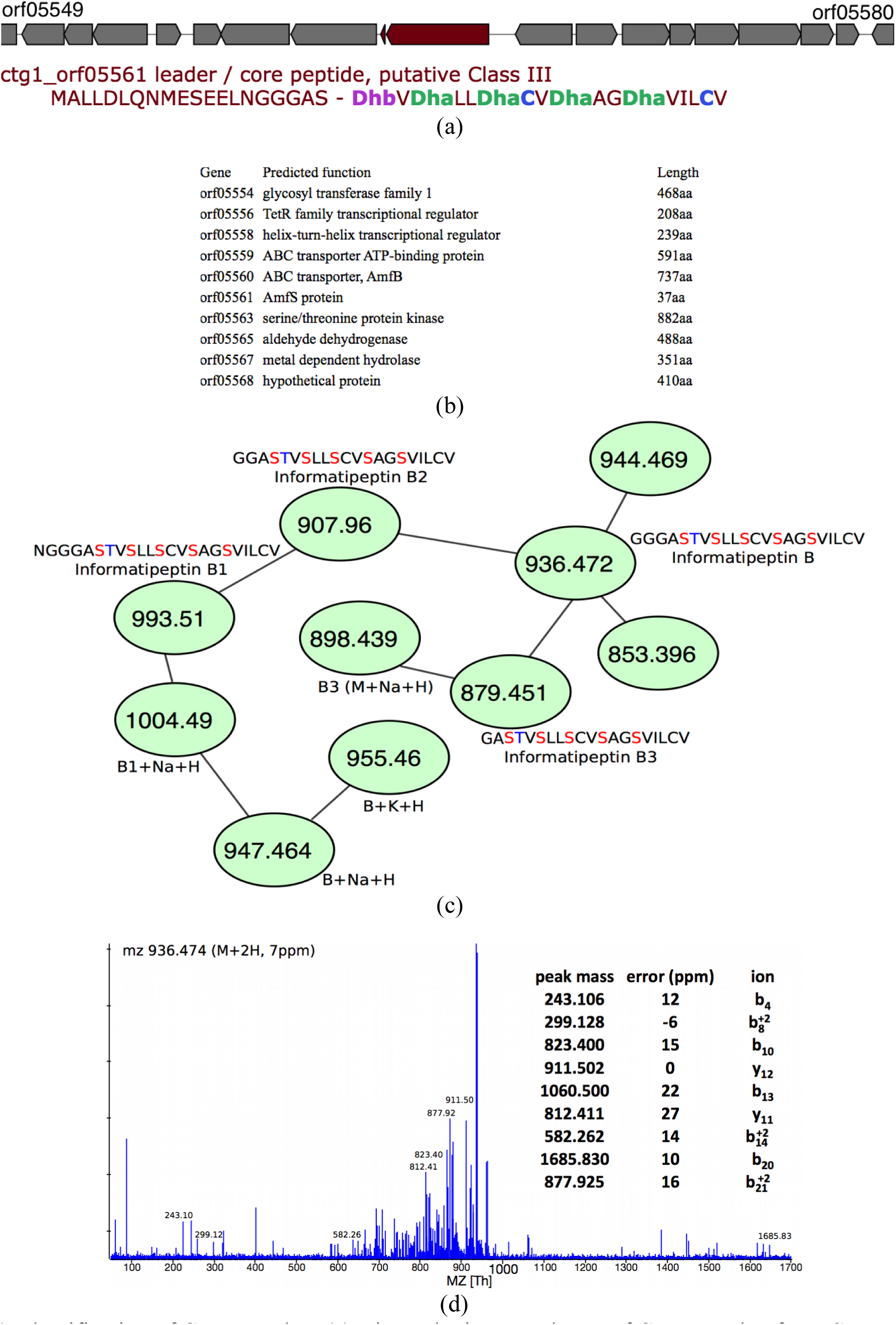
Identification of informatipeptin B. (a) Biosynthetic gene cluster of informatipeptin B from *S. cattleya* and the precursor peptide correctly predicted by antiSMASH. (b) The BGC of informatipeptin B has all essential class III lanthipeptide genes, a lanM-like enzyme, a transporter, and a regulator. (c) Spectral network revealed a plethora of compounds similar to informatipeptin B. Masses shown here are in charge +2 state. Dehydrated serines are shown in red and dehydrated threonines are shown in blue (d) Tandem mass spectrum of informatipeptin B (score 9, p-value 1•10^−12^).

Compound X (DTGGCSGLCTVLVCTVIVC) is a novel lantibiotic that was identified with p-value 6•10^−13^ in the ACTI dataset and that shows no homology to any of the known RiPPs (Figure 4). One of the genes within its BGC shows similarity to genes encoding class I lantibiotics (e.g., subtilin). The most abundant analog of Compound X (mass 1769.80 Da) corresponds to four dehydrations (unmodified peptide mass of 1841.84). The spectral network of Compound X revealed an additional Compound X1 analog (truncation by one amino acid).

**Figure 4.**
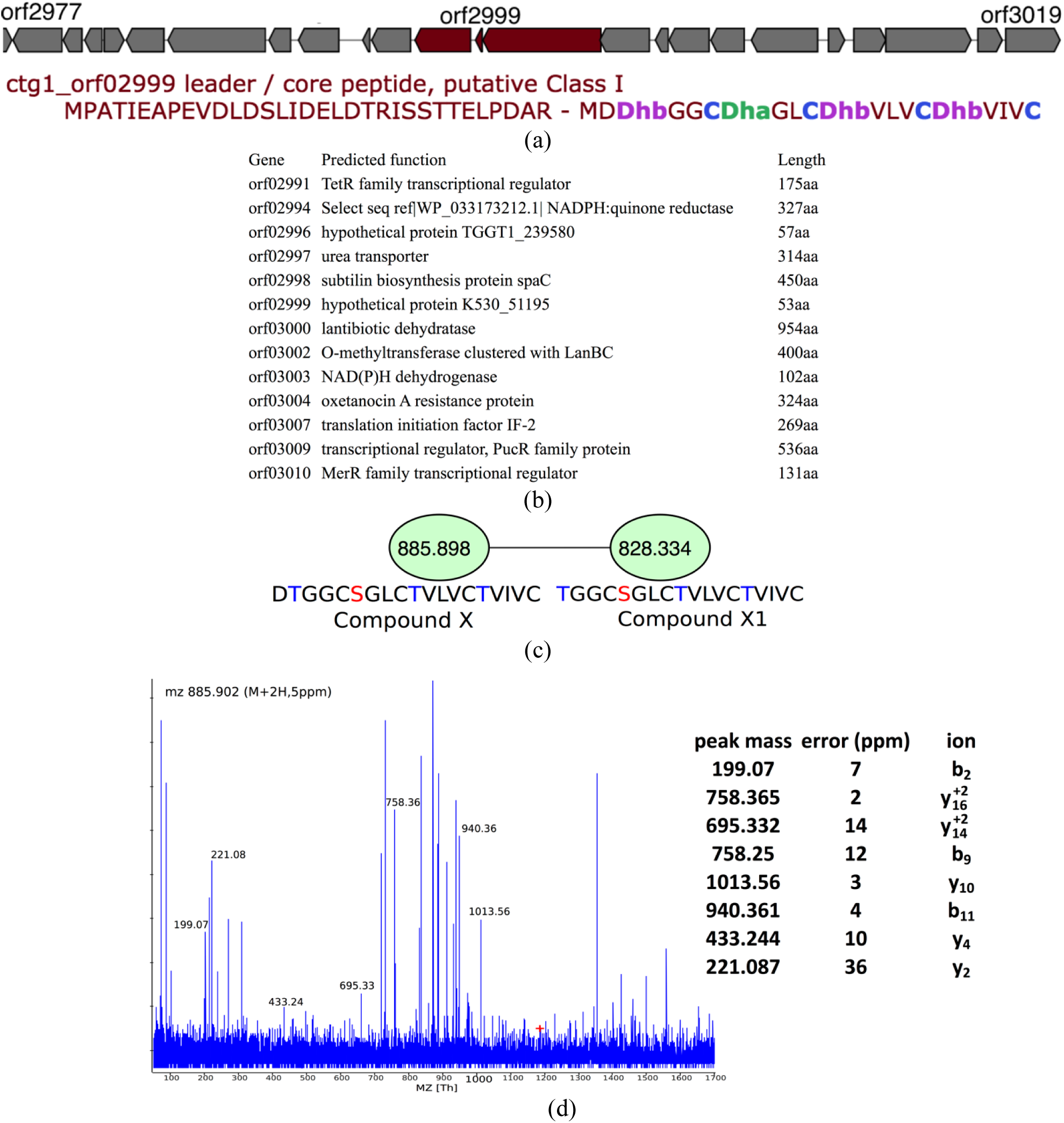
Identification of Compound X. (a) Biosynthetic gene cluster of Compound X from *Streptomyces* sp. and the precursor peptide as correctly predicted by antiSMASH. (b) The gene cluster of Compound X has several genes with *ca*. 60% similarity to genes from BGCs encoding subtilin and other class I lanthipeptide, suggesting that Compound X is a class I lanthipeptide. (c) Spectral networking revealed a analog of Compound X, called Compound X1, lacking the N-terminal aspartic acid residue. Dehydrated serines are shown in red and dehydrated threonines are shown in blue (d) Tandem mass spectrum of Compound X (score 8, p-value 6.0 • 10^−13^).

Compound Y (LLGRHGNDRLILSKN) is a novel lassopeptide that was identified with a p-value of 5•10^−31^ in the ACTI dataset and that shows distant homology to microcin J25 (Figure 5). Two of the genes in its BGC are similar to microcin J25 hypothetical genes McjB and McjC from *S. avertimilis* with a 78% identity for both. The most abundant analog of Compound Y (mass 1645.962 Da) corresponds to a −59 Da modification (unmodified peptide mass of 1704.964 Da).

**Figure 5.**
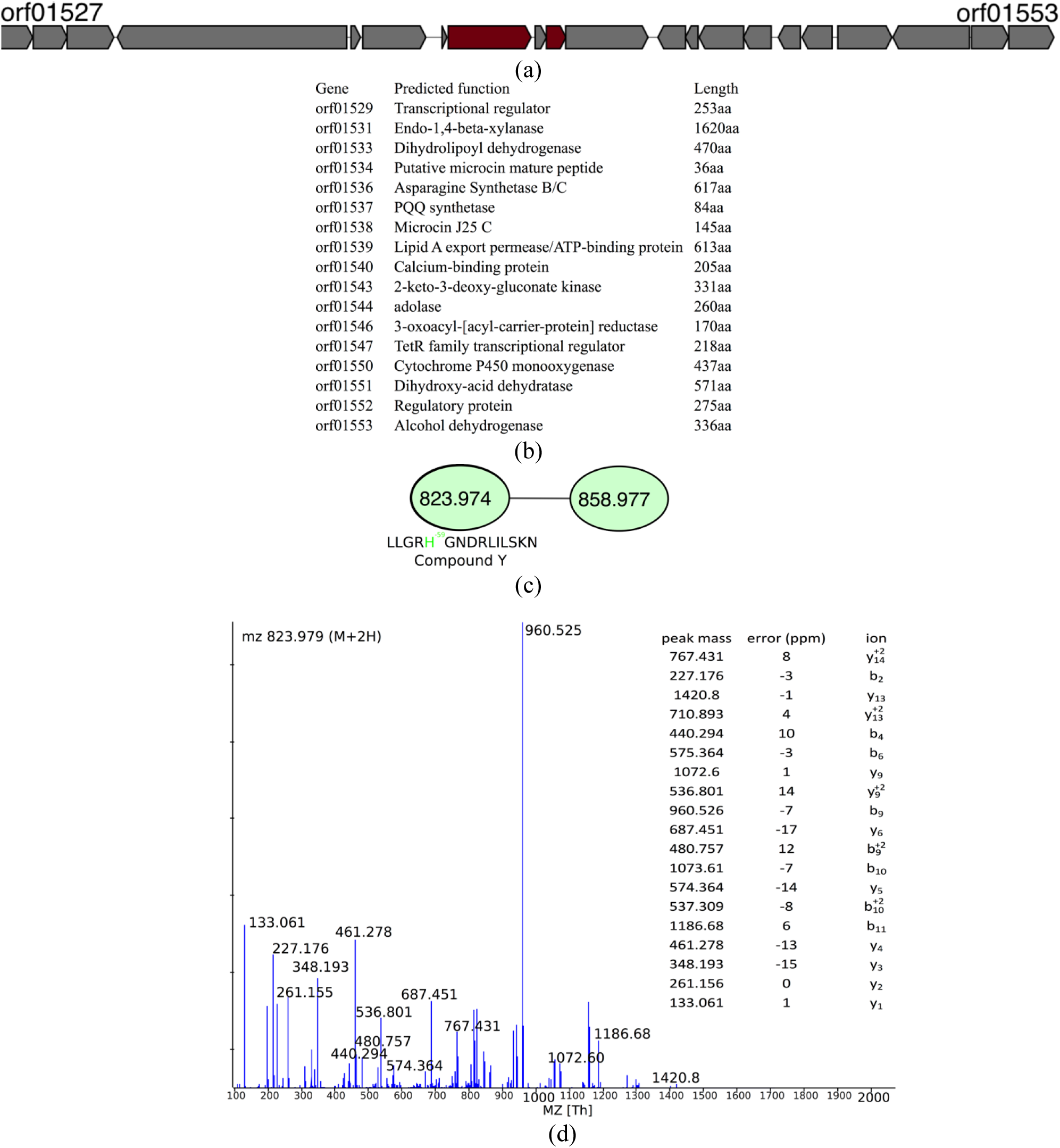
Identification of Compound Y. (a) Biosynthetic gene cluster of Compound Y from *S. viridochromogenes*. AntiSMASH failed to predict any precursor peptide for this gene cluster. (b) The biosynthetic gene cluster of Compound Y has several genes with similarity to the biosynthetic gene cluster of lassopeptide microcin J25. (c) Spectral network of Compound Y (d) Tandem mass spectrum of Compound Y (score 18, p-value 5•10^−31^). MetaRiPPquest discovered Compound Y in blind modification search mode, and assigned a modification of −59Da to the histidine residue, shown in green. While *m/z* 858.97 clusters with Compound Y in the spectral network, MetaRiPPquest does not identify it with a low p-value.

Compound Z (DATITTVTVTSTSIWASTVSNHC) is a new RiPP identified in the SPACE dataset that shows no similarity to any known RiPP. It was not identified in the motif-ORF mode since antiSMASH failed to identify the ORF encoding this RiPP. The gene cluster of Compound Z shows similarity to enterotoxin genes. While enterotoxin gene clusters from *Escherichia coli* are known to produce RiPPs^9,54^, this is the first evidence for production of a RiPP by a *Bacillus* enterotoxin BGC.

The most abundant analog of Compound Z (mass 2109.064 Da) corresponds to eleven dehydrations and a −87Da N-terminal modification (unmodified peptide mass 2394.126 Da). MetaRiPPquest assigned a p-value of 3•10^−25^ to the PSM formed by Compound Z (Figure 6). There are multiple identical ORFs encoding Compound Z in the lantibiotic gene clusters of strains ISSFR-3F (13 copies), S1-R2-T1 (3 copies) and S1-R3-J1 (3 copies). Using spectral networks, MetaRiPPquest detected two less abundant analogs of Compound Z with −1 Da and −15 Da N-terminal modifications, instead of the −87 Da N-terminal modification in the most abundant analog. Moreover, there are analogs corresponding to the sequence DATITTVTVT with five dehydrations and a −87 Da or −15 Da N-terminal modification, and a analog corresponding to the alternative ORF DATITTVTVTSTSIWASTVSNYC in the same lantibiotic gene cluster with a single H to Y mutation.

**Figure 6.**
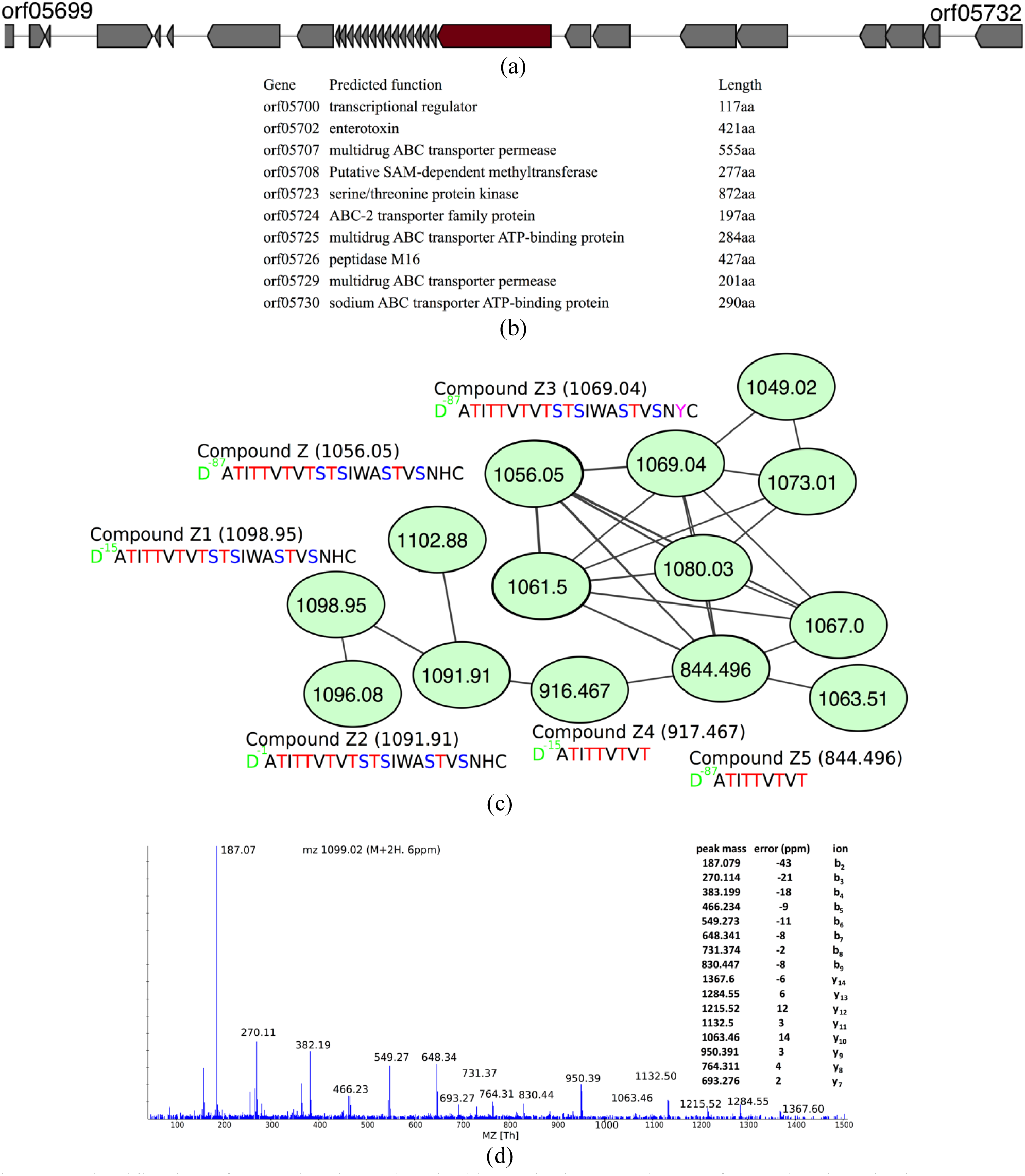
Identification of Compound Z. (a) Biosynthetic gene cluster of Compound Z in *Bacillus* sp. ISSFR-3F. AntiSMASH failed to predict any precursor peptide for this gene cluster. The precursor gene is repeated fourteen times, with thirteen being exactly the same and one differing from the rest by a His to Tyr replacement (b) The BGC of Compound Z (c) Spectral networking revealed analogs of Compound Z with various N-terminal and C-terminal modifications. Dehydrated serines are shown in red and dehydrated threonines are shown in blue (d) The tandem mass spectrum of Compound Z (score 16, p-value 3•10^−25^). MetaRiPPquest discovered compound Z in blind modification search mode, and assigned −87Da, −15Da, and −1Da modifications to the aspartic acid residue, shown in green residue. Compound Z4 and Z5 are discovered in charge +1, while other compounds are charge +2.

Cyanobactin X is a cyclic peptide ISNGYLIP (mass 857.47 Da) with p-value 2•10^−11^ identified in strain PNG22APR06-1 of the CYANO dataset (Figure 7). The cyanobactin X core peptide has no similarity to any of the known cyanobactins. Its BGC, encoded in an 8 kb contig, is missed by antiSMASH. The spectrum of cyanobactin X does not cluster with any other spectra in the spectral network.

**Figure 7.**
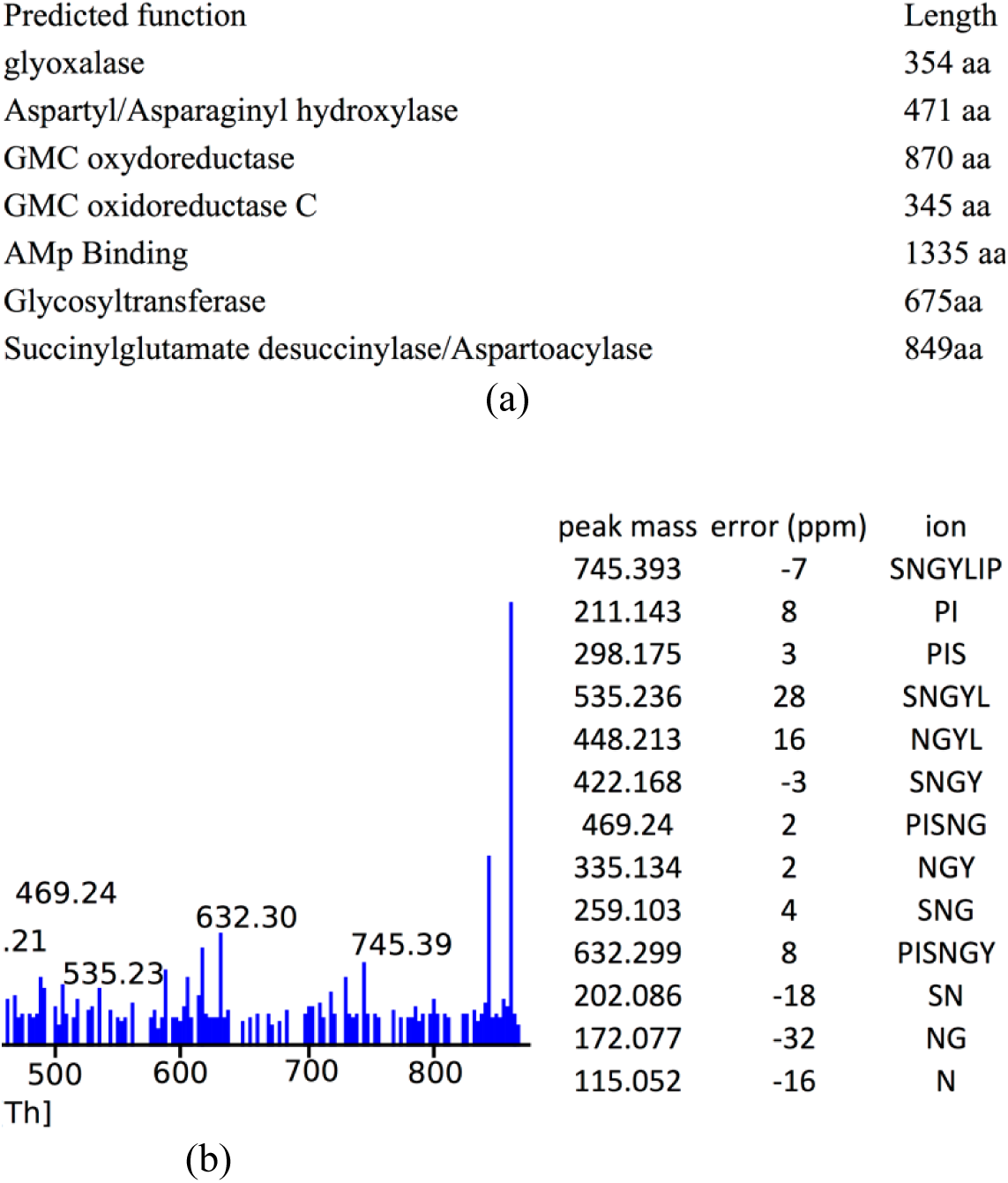
Identification of Cyanobactin X. (a) The biosynthetic gene cluster of cyanobactin X in the Cyanobacterium PNG22APR06-1. AntiSMASH failed to predict this gene cluster. (b) Annotation of the tandem mass spectrum of Cyanobactin X (score 13, p-value 2•10^−17^).

Wewakazole is a cyclic dodecapeptide IS−20APPGVT−20FS−20FP with mass 1140.54 Da, originally discovered in *Lyngbya majuscula* from Papua New Guinea^52^ (S−20 and T−20 stand for oxazole and methyl-oxazole, respectively). MetaRiPPquest identified wewakazole and its BGC with p-value 2•10^−22^ in strain PNG19MAY05-2/7 (Figure 8). AntiSMASH failed to predict any precursor peptide for this gene cluster

**Figure 8:**
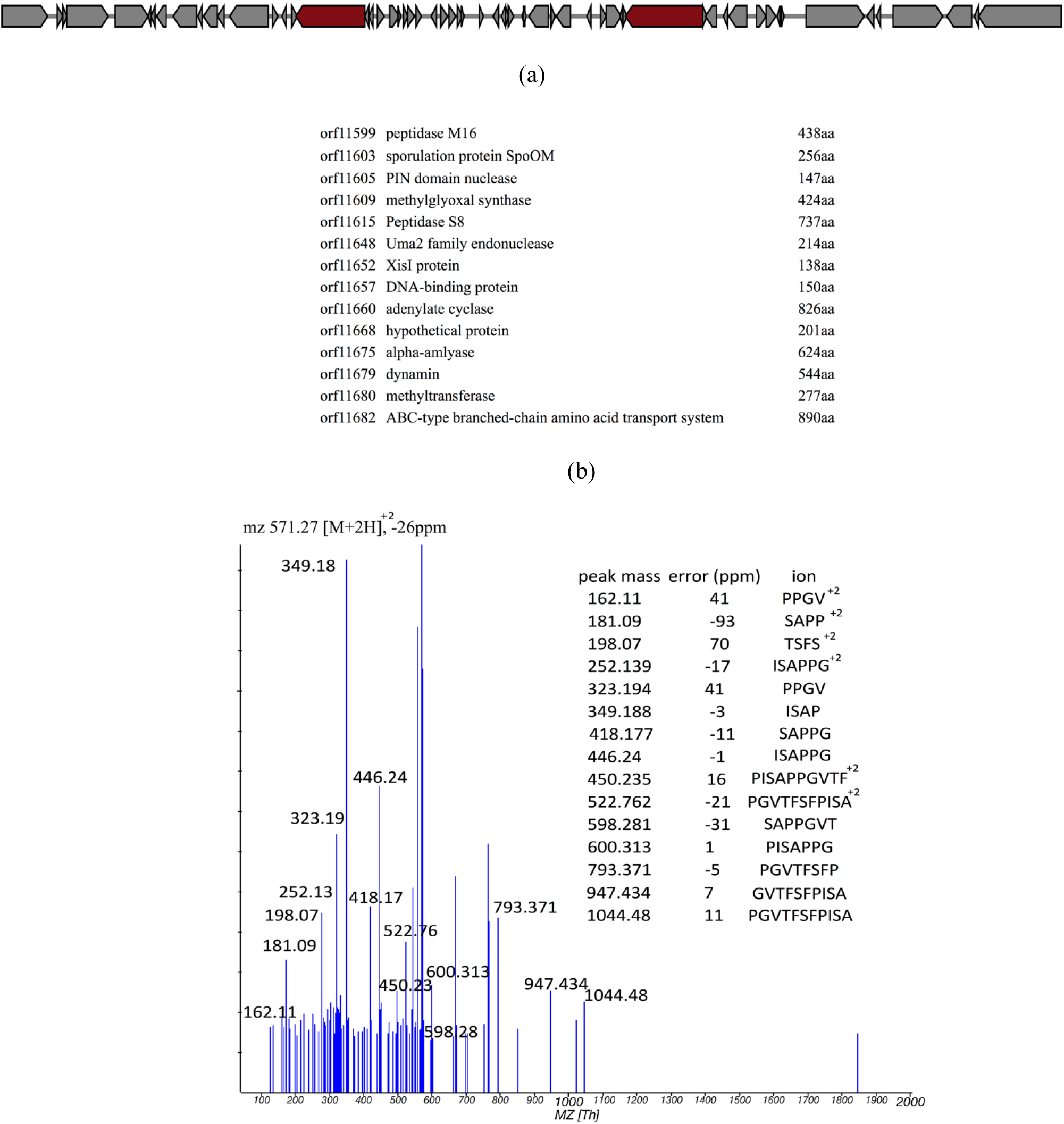
Identification of wewakazole. (a) Biosynthetic gene cluster of wewakazole in the Cyanobacterium PNG19MAY05-2/7. AntiSMASH failed to predict the precursor peptide for this gene cluster by motif search; however, MetaRiPPquest was able to discover this peptide in all-ORF mode. (b) Annotation of the BGC of wewakazole, (c) Annotation of the tandem mass spectrum of wewakazole (score 15, p-value 2•10^−22^.

### Confirmation of wewakazole identification

MetaRiPPquest identified wewakazole in a polar fraction from the extract of strain PNG19MAY05-2/7, a marine cyanobacterium collected at Kape Point, Papua New Guinea. Wewakazole was first reported by one of our groups from another Papua New Guinea collection of *Lyngbya majuscula* (revised to *Moorea producens*) obtained from Wewak Bay^52^. Subsequently a related compound, wewakazole B was isolated from a Red Sea collection of this cyanobacterium^55^. To confirm and to validate the accuracy of MetaRiPPquest to find and identify new compounds from strain PNG19MAY05-2/7, reverse phase C_18_ column chromatography and preparative HPLC separations were successful in the isolation of 31.2 μg of this compound. The compound possessed the same molecular formula as wewakazole, Cs_59_H_72_N_12_O_12_, based on the molecular ion sodium adduct [M+Na]^+^ in the HRESIMS (*m/z* 1163.5282, Supplementary Figure 1). Its chemical identity was further confirmed utilizing ^1^H, HSQC and HMBC NMR data, which allowed for direct comparison with data previously reported for wewakazole (Supplementary Figure 2, 3, 4)^52^. Moreover, the tandem mass spectrum and retention time of the isolated compound matched the data previously reported for wewakazole (Supplementary Figures 5, 6)^52^. Furthermore, the ECCD spectrum resembled that of wewakazole B^53^, and the specific rotation showed the same sign as previously reported for wewakazole^52^, excluding the possibility of an enantiomeric relationship of this isolate to that of wewakazole (Supplementary Figure 7). Thus, the compound identified by MetaRiPPquest was isolated and its identity was confirmed as wewakazole.

### Comparison of extraction/fractionation strategies

MetaRiPPquest can assist in determining the optimal extraction strategy for novel RiPP discovery. As an example, nine strategies were used for fractionation of the samples, and among them strategy H, 25% Methanol and 75% Ethyl acetate, is the only strategy capable of detecting both wewakazole and Cyanobactin X.

## Discussion

While recent genome mining efforts have revealed over 20,000 hypothetical RiPP-encoding BGCs^23^, only 35 RiPPs have been identified that match to these BGCs. To keep pace with the speed of microbial genome sequencing, high-throughput methods for structure elucidation of RiPPs by combining metagenomics, genome mining, and peptidomics are needed. MetaRiPPquest extends our previous RiPPquest tool (limited to lanthipeptides) to lassopeptides, LAPs, linaridins, glycocins, proteusins, and cyanobactins, and enables the blind search for RiPPs with unusual modifications.

Articles describing RiPPs are usually limited to the analysis of a single peptide or a few related peptides. The first application of MetaRiPPquest revealed many known RiPPs, as well as their unknown analogs, and five novel RiPPs (three lantibiotics, one lassopeptide, and one cyanobactin) along with their numerous analogs, from only six spectral datasets. This result provides optimism that MetaRiPPquest can potentially make RiPP identification as robust as peptide identification in traditional proteomics. The increased robustness was validated by the isolation of the RiPP metabolite wewakezole, and its structure was confirmed by orthogonal approaches, confirming that the MetaRiPPquest prediction was correct. In contrast to the existing genome mining approaches that rely on known BGC motifs^28^, MetaRiPPquest in the all-ORF mode has the ability discover new BGCs (with previously unknown motifs) that encode novel RiPPs (e.g. compound Z and cyanobactin X) that are very different from currently known RiPPs and thus are not captured by the existing genome mining tools.

Finally, algorithmic improvements in MetaRiPPquest have resulted in a two orders of magnitude increase in speed compared to RiPPquest, thus enabling searches of the entire GNPS infrastructure against metagenomic information.

## METHODS

### MetaRiPPQuest algorithm

Below we describe various steps of MetaRiPPquest.

#### (a) Selecting a biological sample

MetaRiPPquest works both with DNA sequencing datasets from isolate microbes and bacterial/fungi communities.

#### (b) Generating DNA sequencing data

MetaRiPPquest works with short Illumina reads or pre-assembled genomes/metagenomes.

#### (c) Assembly of DNA sequencing data

MetaRiPPquest assembles reads using SPAdes^26^ or MetaSPAdes^27^.

#### (d) Identifying putative RiPP precursor peptides and constructing a database of putative RiPP structures

MetaRiPPquest uses antiSMASH^28,56^ for genome mining. AntiSMASH identifies putative RiPP-encoding BGCs, searches for known precursor peptide motifs in all ORFs within these BGCs, and constructs the set of putative core peptides. It also identifies modification enzymes in each RiPP-encoding BGC and uses the observation that RiPPs are typically encoded in a 10 kb window centered at a modification enzyme gene.

This approach usually results in a small number of putative core peptides per BGC, resulting in a fast and specific but non-sensitive peptidogenomics approach. As an alternative to this motif-ORF approach, the all-ORF approach (i) starts with all RiPP-encoding BGCs identified by antiSMASH, (ii) constructs a 10 kb windows centered at each modification enzyme of the identified RiPP-encoding BGC, and (iii) identifies all ORFs shorter than a pre-defined threshold (the default is 200 amino acids) as the putative precursor peptide^12^. This approach is sensitive but less specific than the motif-ORF approach.

After all modification enzymes in each RiPP-encoding BGC are identified, MetaRiPPquest considers the corresponding modifications for all suffixes of the identified ORFs within these BGC to construct the target RiPP structure database by considering all possible combinations of modifications (consistent with the modification enzyme) on all possible residues. The decoy RiPP structure database is similarly constructed, starting from decoy ORFs. MetaRiPPquest constructs a decoy database of precursor peptides by random shuffling of the target precursor peptide^57^.

#### (e) Generating tandem mass spectra

Samples were extracted using various fractionations and spectra were collected using various instruments described in supplementary material.

#### (f) Matching spectra against the constructed RiPP structure database

MetaRiPPquest uses a modified version of Dereplicator^29^ for searching spectra against the database of putative RiPP structures as follows (i) theoretical spectra for all peptides in the target and decoy RiPP structure database are constructed, (ii) PSMs are generated and scored, (iii) p-values of PSMs are computed using MS-DPR^58^, (iv) false discovery rates are computed using the decoy database, (v) statistically significant PSMs are output as putative RiPP identifications.

While exhaustive generation of candidate RiPPs and scoring by Dereplicator is feasible when a small number of modifications are considered, the running time rapidly increases with the increase in the number of modifications. We use the spectral alignment technique to efficiently find modifications of the core peptide that best matches the spectrum^59-61,12^. Moreover, to handle blind modifications we adapted the unrestrictive modification search algorithm^61^. This dynamic programming approach restricts the number of modifications and penalizes high score matches with more than one modification.

While the dynamic programming approach from RiPPquest^12^ can handle modifications in linear peptides, it is not applicable to cyclic peptides. MetaRiPPquest uses a brute-force approach to search all RiPP modifications of each candidate cyclic peptide against all spectra. We do not currently perform blind modification searches for cyclic peptides due to the inherent computational complexity.

#### (g) Enlarging the set of identified RiPPs via spectral networking

The set of found RiPPs is enlarged via spectral networks^30,31^.

### Confirmation of wewakazole structure

HESIRMS data was collected using an Agilent 6230 Accurate-Mass TOFMS in positive ion mode by the UCSD Chemistry and Biochemistry Mass Spectrometry Facility. UV-Vis data were recorded on a Beckman Coulter DU 800 spectrophotometer at room temperature in MeOH (λ_max_ at 214 nm and 217 nm). The ECCD spectrum was measured in MeOH using an Aviv 215 CD spectrometer. Optical rotation was measured at 25 °C using a JASCO P-2000 polarimeter ([α]^25^ −3.9 (*c* 0.022, MeOH) (lit.,^52^ [α]^21^_D_ −46.8 (*c* 0.41, MeOH)). A Bruker AVANCE III 600 MHz NMR with a 1.7 mm dual tune TCI cryoprobe was used to record ^1^H, HMBC and HSQC NMR data at 298 ^o^K with standard Bruker pulse sequences. A Varian Vx 500 NMR with a cold probe and z-gradients was used to record ^1^H NMR data at 298 K with standard pulse sequences. NMR data were recorded in CDCh and calibrated using residual solvent peaks (*δ_H_* 7.26 and *δ_c_* 77.16).

For LC-MS analysis, a Thermo Finnigan Surveyor HPLC System was used with a Phenomenex Kinetex 5 μm C18 100 × 4.6 mm column coupled to a Thermo-Finnigan LCQ Advantage Max Mass Spectrometer. Samples were separated using a linear gradient with (A) H_2_O + 0.1% FA to (B) CH_3_CN + 0.1% FA at a flow rate of 0.6 mL/min. The gradient started with a 5 min isocratic step at 30% B followed by an increase to 99% B over 17 min, which was held at 99% B for 5 min and then moved to 30% B in 1 min, and then held for 4 min. Mass spectra were acquired with an ESI source ranging from *m/z* 200-1600.

Preparative HPLC was done using a Kinetex 5 μm C18 150 × 10.0 mm semi-preparative column coupled to a Thermo Dionex Ultimate 3000 pump, RS autosampler, RS diode array detector, and automated fraction collector.

### Isolation of wewakazole

The fraction in which metaRiPPquest identified wewakazole from sample PNG19MAY05-2/7 is identified here as 1648H. This fraction (26.5 mg) was initially separated using a 500 mg/8mL Xpertek^®^ C_18_ SPE cartridge with 100% CH_3_CN to yield 10.7 mg after concentration under vacuum. The compound was isolated from this eluent by semi-preparative HPLC using a linear gradient with (A) H_2_O + 0.1% FA to (B) CH_3_CN at a flow rate of 4 mL/min, and the chromatogram was monitored at 218 nm. The gradient started with a 5 min isocratic step at 40% B followed by an increase to 95% B in 25 min. Approximately 2.5 mg of the sample were injected per run to yield 31.2 μg of wewakazole (*t*_R_=13.0 min).

## Code availability

MetaRiPPquest is available both as a command line tool and as a web application at http://metarippquest.metabologenomics.org.

## Acknowledgement

The work of H.M., P.D. and P.A.P. was supported by NIH 2-P41-GM103484. P.D. is supported by GM097509. A.G., A.M., A.S., A.K. and P.A.P. were supported by Russian Science Foundation 1450-00069. T.L.K., P.D., L.G. and W.H.G. were supported by NIH 2R01GM107550. L.G. and W.H.G. were supported by NIH R01GM118815. JTM was funded by NSF GRFP DGE-1144086. We thank the implementation team of the Microbial Observatory (Microbial Tracking-1) project at NASA Ames Research Center and sample processing/isolation of microbes by Aleksandra Checinska Sielaff, JPL. Part of the research described in this publication was carried out at the Jet Propulsion Laboratory, California Institute of Technology under a contract with NASA. The work of N.K.S. and K.V. was funded by NASA 19-12829-26 and 19-12829-27. C.B.N. was supported by postdoctoral fellowship from NCI/NIH Training Program in the Biochemistry of Growth Regulation and Oncogenesi (T32 CA009523). T.L.K was supported by Vaadia-BARD Postdoctoral Fellowship Award no. FI-494-13. T.L. was supported by CAPES Foundation for Research Fellowship (13425-13-7). The work of I.F. was supported, in part, by NSF awards ABI-1551363 and ABI-1458359.

